# Loss of Peroxisomal Hydroxypyruvate Reductase Inhibits Triose Phosphate Isomerase but Stimulates Cyclic Photosynthetic Electron Flow and the Glc-6P-Phosphate Shunt

**DOI:** 10.1101/278580

**Authors:** Jiying Li, Sarathi M. Weraduwage, Alyssa L. Preiser, Sean E. Weise, Deserah D. Strand, John E. Froehlich, David M. Kramer, Jianping Hu, Thomas D. Sharkey

## Abstract

The oxygenation of ribulose 1,5-bisphosphate by Rubisco is the first step in photorespiration and reduces the efficiency of photosynthesis in C_3_ plants. Our recent data indicates that mutants in photorespiration have increased rates of photosynthetic cyclic electron flow around photosystem I. We investigated mutant lines lacking peroxisomal hydroxypyruvate reductase to determine if there are connections between 2-PG accumulation and cyclic electron flow. We found that 2-PG is a competitive inhibitor of triose phosphate isomerase (TPI), an enzyme in the Calvin-Benson cycle that converts glyceraldehyde 3-phosphate to dihydroxyacetone phosphate. This block in metabolism could be overcome if glyceraldehyde 3-phosphate is exported to the cytosol where the cytosolic triose phosphate isomerase could convert it to dihydroxyacetone phosphate. We found evidence that carbon is reimported as Glc-6P-phosphate forming a cytosolic bypass around the block of stromal TPI. However, this also stimulates a Glc-6P-phosphate shunt, which consumes ATP, which can be compensated by higher rates of cyclic electron flow.

**Once Sentence Summary:** Triose phosphate isomerase is inhibited in plants lacking hydroxypyruvate reductase 1 and this is overcome by exporting triose phosphate to the cytosol and importing Glc-6P, which stimulates a Glc-6P-phosphate shunt and cyclic electron flow.

## INTRODUCTION

Photorespiration occurs when ribulose bisphosphate (RuBP) is oxygenated instead of carboxylated by Rubisco (Bowes et al., 1971). Oxygenation is an unavoidable consequence of the reaction mechanism of Rubisco, and once oxygenation occurs the remainder of photorespiratory metabolism reduces the cost of oxygenation (Andrews and Lorimer, 1978). Photorespiration reduces photosynthesis by three mechanisms: (1) reduction of Rubisco efficiency (increase in apparent *K*_m_ for CO_2_); (2) diversion of ATP and NADPH from carboxylation products to the photorespiratory pathway; and (3) release of CO_2_ in the photorespiratory pathway. These three effects significantly reduce the photosynthetic rate in C_3_ plants (Sharkey, 1988) and likely provided evolutionary pressure that resulted in C4 metabolism (Sage et al., 2012). Photorespiration is also a source of H_2_O_2_ and can affect the redox status of plant cells (Foyer et al., 2009).

Although photorespiration will decline as atmospheric CO_2_ increases, it will remain a significant issue for C_3_ photosynthesis for a long time (Walker et al., 2016b). Using a spreadsheet available here (Sharkey, 2016) it can be calculated that the ratio of oxygenation to carboxylation at 25°C for a plant with Rubisco with kinetics found in *Arabidopsis thaliana* would be 0.53 at preindustrial CO_2_ of 290 ppm, 0.39 at today’s level of ∼400 ppm, and 0.26 when the CO_2_ concentration is 600 ppm (assuming that CO_2_ at Rubisco is 50% of that in air because of diffusion resistances at the stomata and in the mesophyll). Engineering alternative photorespiration pathways to ameliorate the negative effects of photorespiration on plant growth is currently under investigation (Kebeish et al., 2007; Peterhänsel et al., 2013; Dalal et al., 2015; Xin et al., 2015; Betti et al., 2016; Engqvist and Maurino, 2017).

Photorespiratory metabolism converts two molecules of the first product of oxygenation, 2- phosphglycolate (2-PG), to one molecule of 3-phosphoglycerate (PGA) and one molecule of CO_2_. Ordinarily, photorespiration proceeds at the rate needed to metabolize all of the 2-PG produced by oxygenation. Douce and Heldt (2000) conclude that “the only control step in the photorespiratory cycle is the level of competition between O_2_ and CO_2_ for binding to Rubisco.” A number of the enzymes and transporters needed for photorespiration were discovered because plants lacking genes for these proteins can grow in a high CO_2_ atmosphere but not in air (Somerville and Ogren, 1979; Somerville, 1984).

It is assumed that the intermediates of photorespiration reduce the capacity of the Calvin-Benson cycle (Betti et al., 2016). Anderson (1971) showed that a very low concentration of 2-PG significantly inhibited triose phosphate isomerase (TPI), an enzyme that converts glyceraldehyde 3-phosphate (GAP) to dihydroxyacetone phosphate (DHAP). Somerville and Ogren (1979) showed that plants lacking phosphoglycolate phosphatase accumulate 2-PG and, based on Anderson’s work, proposed this would cause more triose phosphate to be exported from chloroplasts, favoring sucrose synthesis. By altering the amount of phosphoglycolate phosphatase, Flügel et al. (2017) confirmed the inhibition of TPI by 2-PG and found less starch in plants with high 2-PG but did not find increased sucrose. Xu et al. (2009) reported that plants lacking some activity of glycolate oxidase had reduced activation of Rubisco and less mRNA for Rubisco activase than controls when grown in air but more Rubisco activase mRNA when grown in elevated CO_2_ (0.5%). Walker et al. (2016a) found reduced Rubisco activation ratio in plants lacking the glycerate/glycolate transporter PLGG1. Flügge et al. (1980) showed that glyoxylate could cause acidification of the stroma and so inhibit pH-sensitive enzymes in the Calvin-Benson cycle, but this effect was not specific to glyoxylate, other organic acids had the same effect. Kelly and Latzko (1976) reported that 2-PG (but not glycolate) inhibited phosphofructokinase, which they postulated could interfere with starch breakdown at night. However, there should be no 2-PG present when starch is breaking down and current thinking about starch breakdown does not include a role for phosphofructokinase (Stitt and Zeeman, 2012).

Most of the enzymes and transporters of photorespiratory metabolism are essential; plants that lack them are very stunted or die when grown in air. However, plants missing NADH-dependent hydroxypyruvate reductase in the peroxisome (*hpr1*) show only modest effects (Timm et al., 2008; Cousins et al., 2011; Timm et al., 2012). It is known that hydroxypyruvate can leave the peroxisome and be acted on by a cytosolic HPR (encoded by *HPR2*) that prefers NADPH (Kleczkowski et al., 1988; Timm et al., 2008). In addition, there is a multifunctional glyoxylate/hydroxypyruvate reductase in the chloroplast that prefers NADPH (Tolbert et al., 1970) that has now been identified as HPR3 (Timm et al., 2011). Loss of *HPR1* and *2*, or all three, results in severe growth defects in air but not high CO_2_ (Timm et al., 2008; Cousins et al., 2011; Timm et al., 2011). In addition, it has been hypothesized that some carbon leaves the photorespiratory pathway as glycine or serine (Harley and Sharkey, 1991; Busch et al., 2018). Despite these alternatives, plants lacking *HPR1* are typically stunted indicating that reduction of hydroxypyruvate is necessary for optimal photosynthesis.

In a recent survey of mutants of peroxisomal proteins using the Dynamic Environment Phenotype Imager (Cruz et al., 2016), many of the mutants exhibited significant phenotypes under high and fluctuating light but little phenotype under the low, constant light normally used to grow *Arabidopsis thaliana*. For instance, *hpr1-1, plgg1*, and *cat2-1* showed a significant increase in cyclic electron flow around photosystem I (CEF).

In this study, we investigated potential mechanisms by which a block in photorespiration could stimulate CEF. HPR1 was chosen as the focus of the study because *hpr1* plants grow reasonably well but still show a strong stimulation of CEF. Also, HPR1 was identified as playing a role in drought tolerance (Li and Hu, 2015).We measured amounts of accumulated 2-PG. We then looked at how 2-PG might affect the Calvin-Benson cycle and how those effects might lead to CEF. We postulate that the block of stromal TPI can be bypassed by export of GAP followed by reimport of carbon into the chloroplast. However, this cytosolic bypass of the gluconeogenic reactions of the Calvin-Benson cycle leads to a stimulation of a Glc-6-P shunt (Sharkey and Weise, 2016) that consumes ATP, leading to CEF.

## RESULTS

### Growth, Chlorophyll, Carotenoid Contents and Rubisco Activity of *hpr1* Plants

Both *hpr1-1* and *hpr1-2* plants showed a stunted phenotype when grown in air in these experiemnts (Fig. 1 A). In addition to being smaller, the mutants had significantly less chlorophyll (Fig. 1 B) and carotenoid (Fig. 1 C) than the wild type (WT) Col-0. On the other hand, the total extractable activity of Rubisco was the same in all three lines (Fig. 2 A). The amount of Rubisco activase protein was less in the two mutant lines than WT (Fig. 2 B). The activation state of Rubisco was modestly lower in low light (125 µmol m^−2^ s^−1^) in *hpr1-1* but not *hpr1-2* (Fig. 2 C). After incubation in high light (1000 µmol photons m^−2^ s^−1^) the activation state of Rubisco was higher in all three lines but did not differ among lines (Fig. 2 D).

**Figure 1:**
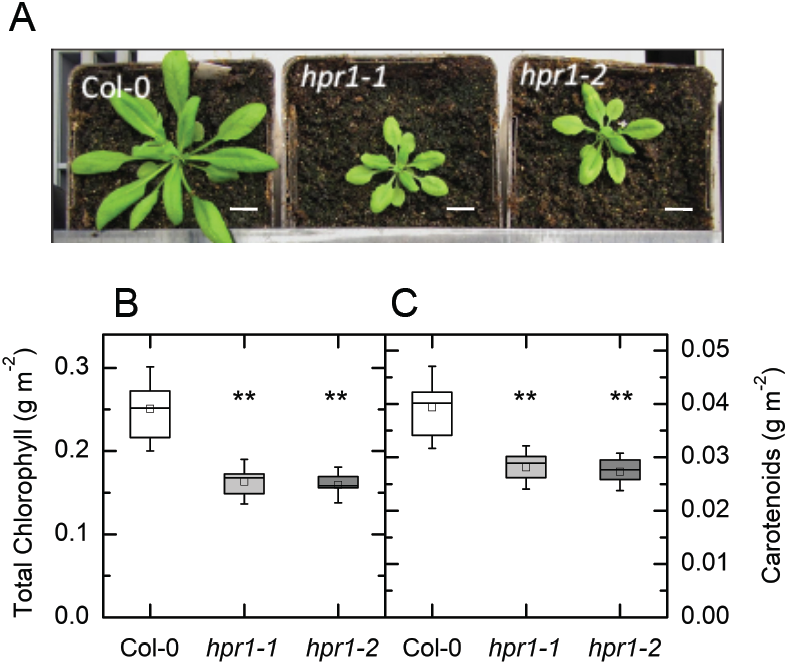
Rosette appearance of WT, *hpr1-1* and *hpr1-2* plants and pigments. **A)** Pictures of two examples of each of three lines of *Arabidopsis thaliana* growing in soil for four weeks. **B)** Chlorophyll and **C)** carotenoid contents. Chlorophyll and carotenoids were measured as described in the materials and methods on leaves of four-week-old plants. White bar in A) is = to 0.5 inch. n=6

**Figure 2:**
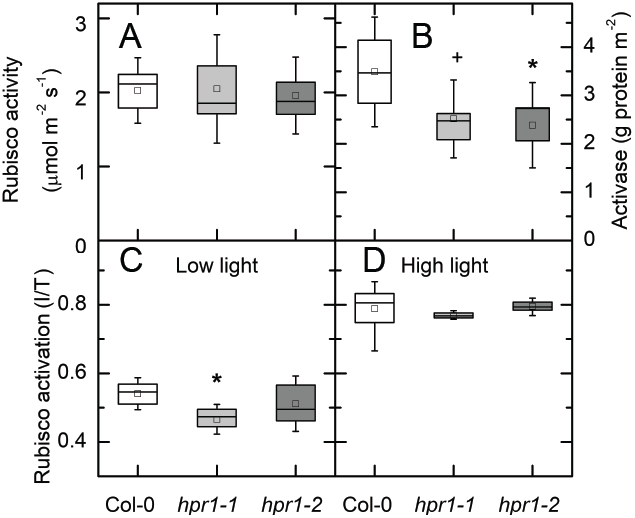
Rubisco activity and activase protein. Total Rubisco activity (**A**) was the same in Col-0, *hpr1-1* and *hpr1-2*. Activase protein (**B**) was measured using antibodies antibodies raised against rubisco activase and a WES capillary electrophoresis instrument from Protein Simple. Rubisco activation state is presented as the ratio between initial (measured as quickly as possible after extraction) and total (after incubation with HCO_3_ and Mg) at 125 (**C**) or 1000 (**D**) µmol photons m^−2^ s^−1^. For a, c, and d n=4-7. For b, data were not different between the low and high light treatment and so was combined, n=7-8.

### 2-PG Content of Leaves is High in *hpr1-1* and *hpr1-2*

The content of 2-PG was measured by LC/MS-MS. The results are expressed as peak area for 2-PG relative to the peak area of the internal standard, (^13^C_6_ fructose 1,6-bisphosphate). The 2-PG content was significantly increased in the *hpr1-1* plants (Fig. 3). In *hpr1-2* plants the level appeared higher (Fig. 3) but the difference did not reach statistical significance.

**Figure 3:**
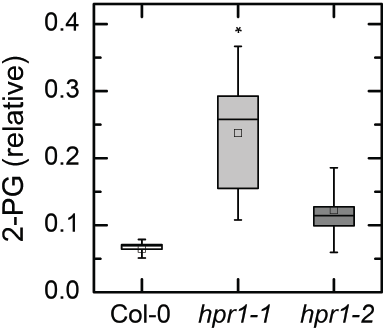
2-phosphoglycolate (2-PG) amounts in leaves of Col-0 and two HPR mutants. 2- PG was measured by HPLC-MS/MS leaves were harvested after 6 hr in 125 or 1000 µmol photons m^−2^ s^−1^. Because the data for both light intensities was indistinguishable they were combined to increase statistical power. Data are expressed as peak area of 2-PG divided by peak area of the internal standard relative to weight of plant material extracted. The results in high and low light were not different and so all of the data was combined. n=7-8

### Effect of 2-PG on TPI Activity

The effect of 2-PG on TPI was tested. 2-PG was a strong inhibitor of TPI and was competitive with GAP (Fig. 4). The *K*_*i*_ was 24.2 µM, between the two values reported previously (Anderson, 1971; Flügel et al., 2017). The relative effect this would have on TPI depends on the concentrations of GAP and 2-PG. To test whether the 2-PG was limiting the activity of TPI in vivo we measured the concentrations of the triose phosphates. The concentration of DHAP was the same in Col-0 and *hpr1-1* but the concentration of GAP was about two-fold higher in *hpr1-1* than in Col-0 (Fig. 5) (note that GAP data have been multiplied by 10). Because GAP is the molecule first produced in the stroma, an inhibition of TPI would be expected to cause a buildup of GAP. At equilibrium, the DHAP/GAP ratio is expected to be 21.5 (Sharkey and Weise, 2012). In Col-0 this ratio was 11.3. and in *hpr1-1* the ratio was 6.1 consistent with a block in TPI activity affecting the Calvin-Benson cycle but also consistent with lack of equilibrium in Col-0.

**Figure 4:**
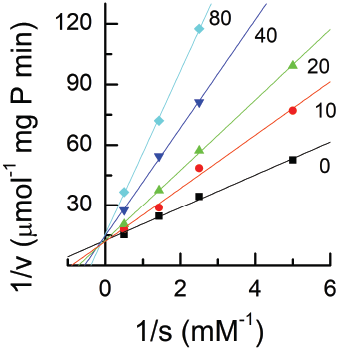
Lineweaver-Burke plot of triose phosphate isomerase activity. Triose phosphate isomerase in crude extracts of Arabidopsis leaves was assayed in the presence of a range of GAP and 2-PG concentrations. The numbers on each line is the concentration of 2-PG in µM. The crossover of the lines at the y axis indicates that 2-PG is a competitive inhibitor.

**Figure 5:**
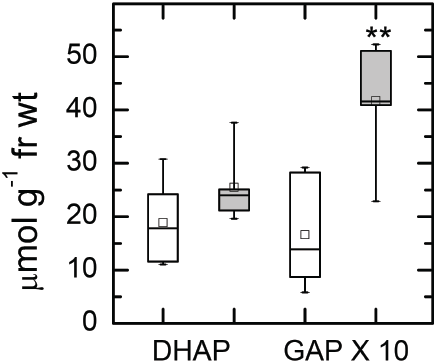
Amounts of DHAP and GAP. Leaves were treated with 1000 µmol photons m^−2^ s^−1^ for six hours, then clamped with liquid-nitrogen-cooled copper blocks to stop metabolism in the light. Leaf samples were extracted with perchloric acid, neutralized, and then measured using sequential additions of GAP dehydrogenase and TPI while following NADH production photometrically. Unfilled boxes – 125 µmol photons m^−2^ s^−1^ low light, grey boxes - 1000 µmol photons m^−2^ s^−1^ high light. n=5

### Expression of the GPT2 Gene

It has been hypothesized that plants that lack chloroplastic fructose bisphosphatase might create a bypass by exporting more carbon to the cytosol and then reimporting carbon, possibly as Glc-6-P though the GPT2 transporter (Kossmann et al., 1994; Sharkey and Weise, 2016). We tested whether the *hpr1-1* plants with inhibited stromal TPI could make use of cytosolic TPI (since there would be no 2-PG in the cytosol) by reimporting carbon as Glc-6-P through the GPT2 transporter. When plants were grown and sampled in low light (125 µmol photons m^−2^ s^−1^) there was no difference in the transcript levels for *GPT2* (Fig. 6 A). However, after exposing plants to 1000 µmol photons m^−2^ s^−1^ for 6 hours, transcripts for *GPT2* were up four-fold in Col-0 plants and ten-fold in *hpr1-1* plants (Fig. 6 B) (note the ten-fold difference in the scale). Plants lacking the glycerate/glyoxylate transporter (*plgg1*) also had increased *GPT2* expression. These plants have increased glycolate (Pick et al., 2013) and presumably 2-PG, as found in the *hpr1-1* mutant. However, plants lacking either of two other photorespiratory genes, *cat2*, or *gox1*, did not exhibit increased levels of *GPT2* mRNA beyond that seen in Col-0 (Fig. 6).

**Figure 6:**
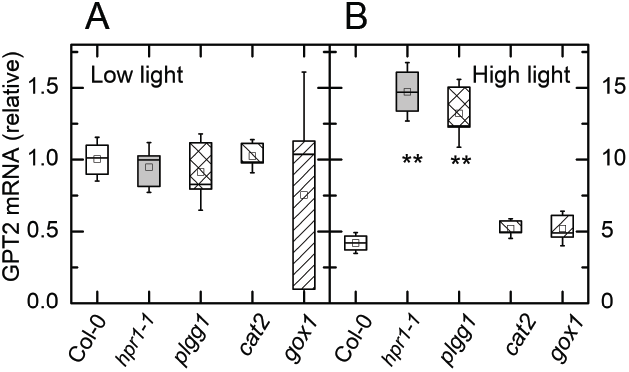
Quantitative RT-PCR analysis of GPT2 transcripts in low and high light. Transcripts were measured in four photorespiratory mutant lines and WT (Col-0) in plants taken from **A**) low light intensity (125 µmol m^−2^ s^−1^) and after six hr **B**) high light (500 µmol m^−2^ s^−1^). Note the scale is 10-fold higher for the high light data, thus GPT2 transcript was about four-fold higher after the high light treatment. n=3

## DISCUSSION

Inhibition of photorespiratory metabolism, other than Rubisco oxygenase activity, results in reduced plant health. A major cause is reduced activity of TPI because of competitive inhibition by 2-PG. Flügel et al. (2017) showed that this led to reduced fructose 1,6-bisphosphate and also sedoheptulose 1,7- bisphosphate, two metabolites that require DHAP for synthesis. Although 2-PG is near the beginning of the pathway and HPR is near the end, loss of HPR activity caused a buildup of 2-PG. Plants lacking HPR1 also accumulate glycolate (Timm et al., 2012). In this study, we found that the ratio of DHAP to GAP was reduced by nearly 50% in *hpr1-1* compared to Col-0. The triose phosphates were not in equilibrium even in the wild type Col-0. A lack of equilibrium at this step was hypothesized in the original publication of the Calvin-Benson cycle (Bassham et al., 1954). They suggested that a lack of equilibrium in this step could explain their observed lower-than-predicted labeling of carbon 4 in sedoheptulose bisphosphate during very short labeling. The lack of equilibrium at TPI can also explain the “Gibbs effect” in which C4 of hexoses is more heavily labeled than C3 (Kandler and Gibbs, 1956; Gibbs and Kandler, 1957). The larger amount of DHAP relative to GAP can also contribute to the disequilibrium of label in carbons 3 and 4 (Ebenhöh and Spelberg, 2018) but the TPI disequilibrium significantly exacerbates the disequilibrium. Therefore, despite its reputation for efficiency, TPI activity is not in excess during photosynthesis and the inhibition of TPI by 2-PG can have a significant effect on metabolism. To overcome the inhibition of TPI, GAP can be exported to the cytosol where, in the absence of 2-PG, TPI can function.

We also tested whether Rubisco activase could play a role in the inhibition of photosynthesis when photorespiratory metabolism is blocked. The growth conditions used in our study resulted in smaller plants with less chlorophyll and carotenoid per leaf area (Fig. 1) and hence the block in photorespiration was affecting plant growth. However, the total activity of Rubisco was the same in Col-0 and two mutant lines (Fig. 2 A). The amount of Rubisco activase protein was less in the mutant lines (Fig. 2 B) consistent with the observation of reduced Rubisco activase message seen by Xu et al. (2009) but Rubisco activation state was lower in only one line and only at low light (Fig. 2 C). Although this could affect how well plants could cope with a stochastic light environment (Taylor and Long, 2017), we conclude from these results that effects on Rubisco activation may be secondary to the main effects of TPI inhibition and the rest of the discussion will focus on the possible consequences of TPI inhibition.

Export of carbon for processing in the cytosol, as proposed here for TPI inhibition, has been proposed for other mutants. Sharkey and Weise (2017) hypothesized that a block in stromal FBPase can be bypassed by export of carbon to the cytosol. Carbon is then reimported into the chloroplast beyond the FBPase step in the Calvin-Benson cycle (Kossmann et al., 1994; Sharkey and Weise, 2016). A mutant lacking fructose 1,6 bisphosphate aldolase also exhibits high rates of CEF (Gotoh et al., 2010). In this mutant carbon could be exported as triose phosphates and then reimported as Glc-6-P to bypass the FBP aldolase step. We hypothesize that in these three cases (and perhaps others) carbon can be exported to the cytosol to bypass missing or limited enzymes of the Calvin-Benson cycle and then reimported into the chloroplast.

However, the gradient for triose phosphate reimport is unfavorable. The phosphate concentration is much higher in the cytosol than in the stroma (Sharkey and Vanderveer, 1989), which aids in triose phosphate export but would slow triose phosphate reimport. On the other hand, the gradient for Glc-6-P is highly favorable for import into the chloroplast (Gerhardt et al., 1987; Sharkey and Vassey, 1989; Szecowka et al., 2013) and could overcome the unfavorable phosphate gradient. Stimulation of *GPT2* expression would allow carbon to be reimported into the chloroplast as Glc-6-P. Transcripts for *GPT2* were elevated in *hpr1-1* and *plgg1* (Fig. 7) but not in *cat2* and *gox1*. We saw that 2-PG was elevated in *hpr1-1*. Pick et al. (2013) saw that glycolate is elevated in *plgg1* and it is very likely that 2-PG is also elevated in this mutant. Since *cat2* and *gox1* are important for redox control but not carbon metabolism, and because there is redundancy of both catalase and *GOX* genes, perhaps *cat2* and *gox1* do not have elevated 2-PG [*cat2* does not have elevated glycolate (Kerchev et al., 2016)] and so do not result in a signal to increase the expression of *GPT2*.

**Figure 7:**
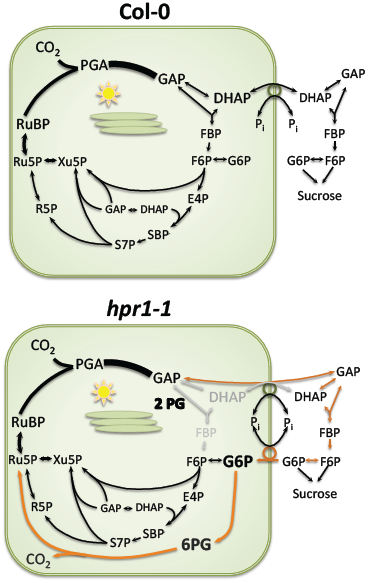
Hypothesized carbon flow in Col-0 and *hpr1-1*. The canonical path of carbon in the Calvin-Benson cycle plus sucrose synthesis is shown for Col-0. In *hpr1-1* plants we hypothesize that 2-PG inhibits TIM forcing export of GAP. Once acted on by cytosolic TIM some DHAP will be reimported but the gradient in phosphate is not favorable for this flux. The triose phosphates in the cytosol will be converted eventually to Glc-6-P. Some unknown signal results in the expression of GPT2 allowing reimport of carbon as Glc-6-P. This allows some Calvin-Benson cycle activity but also stimulates the Glc-6-P shunt (shown in orange). The stimulated Glc-6-P shunt is balanced in terms of carbon and NADPH but reduces ATP availability. Another unknown signal results in cyclic electron flow. PGA = phosphoglyceric acid, GAP = glyceraldehyde 3-phosphate, DHAP, dihydroxyacetone phosphate, FBP = fructose 1,6-bisphosphate, F6P = fructose 6-phosphate, E4P = erythrose 4-phosphate, S7P = sedoheptulose 7-phosphate, R5P = ribose 5-phosphate, XuBP = xylulose 5-phosphate, Ru5P = ribulose 5-phosphate, RuBP = ribulose 1,5-bisphosphate, SBP = sedoheptulose 1,7- bisphosphate, 6PG = 6-phosphoglucanate.

Figure 7 shows a model for the metabolic changes in *hpr1-1*. In Col-0 (top) carbon is exported from the chloroplast for sucrose synthesis but once cytosolic FBPase acts, the carbon can no longer return to the chloroplast because there is only minimal capacity for G1P import (Fettke et al., 2011) and normally no capacity for Glc-6-P import. However, in the *hpr1-1* mutant (bottom) 2-PG inhibits TPI. GAP can be exported and DHAP is made in the cytosol because 2-PG is restricted to the chloroplast. However, phosphate is in high concentration in the cytosol and much lower in the chloroplast (Sharkey and Vanderveer, 1989) making it difficult for DHAP to be imported into the stroma at sufficiently high rates. Moreover, the high concentrations of triose phosphates in the cytosol could lead to FBP synthesis and dephosphorylation. Once FBP is converted to Fru-6-P by cytosolic FBPase, carbon could only reenter the stroma as hexose phosphate. Induction of GPT2 would allow carbon to be reimported as Glc-6-P. Once Glc-6-P is reimported into the chloroplast it can rejoin the Calvin-Benson cycle when it is converted to Fru-6-P by PGI. This would result in a cytosolic bypass around the inhibited TPI.

However, stromal PGI is kinetically limited (Backhausen et al., 1997). The Fru-6-P to Glc-6-P ratios are normally 1:1 in the stroma, well below the predicted ratio and well below the ratio found in the cytosol (Gerhardt et al., 1987; Sharkey and Vassey, 1989; Szecowka et al., 2013). The *K*_*m*_ of PGI for Glc-6-P is much higher than the *K*_*m*_ for Fru-6-P (Schnarrenberger and Oeser, 1974)(Table 1 of this paper shows the opposite but this is in error as can be deduced from the text and abstract and as confirmed by our own unpublished results). GPT2 would allow a very high concentration of Glc-6-P in the stroma. Because stromal Glc-6-P dehydrogenase (G6PDH) has a high *K*_*m*_ when it is reduced (Scheibe et al., 1989; Hauschild, 2003), a high concentration of Glc-6-P could overcome the light-induced changes in G6PDH that normally restrict its activity during the day. This would stimulate the oxidative branch of the pentose phosphate pathway making a Glc-6-P shunt around the Calvin-Benson cycle (Sharkey and Weise, 2016).

The Glc-6-P shunt results in a release of CO_2_. This release would appear similar to the release of CO_2_ during photorespiration. Cousins et al. (2011) reported that *hpr1-1* mutants exhibited higher than expected CO_2_ release that was oxygen dependent. Plants lacking peroxisomal malate dehydrogenase, needed for HPR1 function, also released more than expected amounts of CO_2_ (Cousins et al., 2008). We speculate that the excess CO_2_ release seen in these studies results from the CO_2_ released during the Glc-6-P shunt.

The Glc-6-P shunt consumes 3 ATP but is balanced for NADPH and carbon. Stimulation of this shunt would require significantly more ATP and so CEF could be favored. This provides a mechanism for connecting carbon metabolism changes with electron transport changes. If true, then other photorespiratory mutants that had been shown to contain elevated 2-PG levels should also show elevated CEF. Other mutants that reduce stromal FBP should also show elevated CEF because carbon would avoid these blockages through the cytosolic bypass leading to increased rates of the Glc-6-P shunt. Gotoh et al. (2010) reported a strong stimulation of CEF in a mutant lacking stromal Fru-1,6-bisP aldolase, and plants lacking stromal FBPase also have high rates of cyclic electron flow (Livingston et al., 2010), which is consistent with a cytosolic bypass leading to a Glc-6-P shunt (Fig. 7). There remain unanswered questions, especially how the use of ATP in the Glc-6-P shunt results in a signal to increase CEF and the role of H_2_O_2_ in CEF (Strand et al., 2015) and other metabolism.

Finally, hydoxypyruvate reductase is only required if a significant amount of carbon that originates as 2- PG completes the photorespiratory pathway and becomes glycerate or CO_2_. Some hydroxypyruvate can be reduced by other HPRs (Tolbert et al., 1970; Kleczkowski et al., 1988; Timm et al., 2008; Cousins et al., 2011; Timm et al., 2011). In addition, there are indications that some carbon can leave the photorespiratory cycle as glycine or serine, but that would be limited by how fast nitrogen is made available (Harley and Sharkey, 1991; Busch et al., 2018). While the effect of the loss of HPR1 alone is not as great as the loss of other photorespiratory enzymes, for example 2-PG phosphatase (Somerville and Ogren, 1979), it nevertheless can impair plant growth indicating that a significant fraction of carbon that begins the photorespiratory cycle as a result of oxygenation is reduced by HPR1 and completes the cycle. Nevertheless, because of the alternative paths for hydroxypyruvate reduction, *hpr1* plants are useful for study of the effects of an impairment, but not loss, of photorespiration.

## MATERIALS AND METHODS

### Plant Material

Col-0, *hpr1-1*, and *hpr1-*2 were used. The hpr1 plants (Salk 067724 and 143584) were the same as reported on by (Timm et al., 2008), who showed that no HPR protein was detectable. Plants were grown in Redi-earth potting medium under a 16-h photoperiod. Growth chamber conditions were set to: 120 µmol m^−2^ s^−1^ light intensity, day- and night-time temperatures of 22-23°C and 20°C, and 60% relative humidity. For high light treatment, plants were exposed to a light intensity of 1000 µmol m^−2^ s^−1^ for 6 hr unless otherwise noted.

### Measurement of Chlorophyll and Carotenoid

For chlorophyll measurement, two leaf discs were harvested from 4-week old rosettes, and chlorophyll was extracted in 2 mL of 96% ethanol using a protocol modified from Lichtenthaler and Wellburn (1983). The homogenate was maintained under dark conditions to prevent chlorophyll degradation. Absorbance of each tube at 470 nm, 649 nm and 665 nm were measured with a spectrophotometer. The chlorophyll a, b, and carotenoid content was calculated as follows: chlorophyll a (µg/mL) = 13.95*A*665 - 6.88*A*649; chlorophyll b = 24.96*A*649 - 7.32*A*665; carotenoids = (l000*A*470 - 2.05C_a_ - 114.8C_b_)/245 (Lichtenthaler and Wellburn 1983). Pigment concentration was expressed in g m^−2^ of leaf tissue.

### Measurement of Rubisco Activity and Protein Levels of Rubisco Activase

Leaves from 4-week old rosettes were harvested quickly into liquid N_2_, and stored in a −80 °C freezer. Rubisco activity was measured using a protocol modified from Sharkey et al. (1986) and Sharkey et al. (2001). Leaf tissue was ground with the aid of a Retch mill (Retsch, http://www.retsch.com/) and protein was extracted into 2 mL of extraction buffer (50 mM EPPS pH 8.0, 30 mM NaCl, 10 mM mannitol, 5 mM MgCl_2_, 2 mM EDTA, 5 mM DTT, 0.5% Triton x-100, 1% PVPP, 0.5% casein, 1% protease inhibitor cocktail-P9599 Sigma). To obtain initial activity, 20 µL of extract was added to 80 µL of assay buffer (50 mM EPPS pH 8.0, 5 mM MgCl_2_, 0.2 mM EDTA, 0.5 mM RuBP, 15 mM H^14^CO_3_^−^) in a 7-mL scintillation vial. After a 1 min incubation, 100 µL of 1 M formic acid was added to stop the reaction and release unreacted bicarbonate. The mixture was then dried and the amount of radioactivity that was fixed was determined with the aid of a liquid scintillation analyzer (Tri-Carb2800TR, PerkinElmer). In order to determine total activity, 200 µL of the initial extract was added to 20 µL of activating solution (to give final concentrations of 20 mM MgCl_2_, 15 mM H^14^CO_3_^−^, 61 µM 6-phospho-gluconate) and incubated for 10 min. Total Rubisco activity of the activated sample was assayed as explained above. Each day, radioactivity in 10 µL of the assay buffer was counted to determine specific activity. Based on 1 mCi = 2.22 x 10^9^ distintegrations min^−1^, initial and total Rubisco activity was calculated and expressed as µmol m^−2^ s^−1^. The rates were divided by 0.943 to account for the discrimination against ^14^C by (Roeske and O’Leary, 1984).

To determine Rubisco activase protein content, total protein was extracted using a PlantTotal Protein Extraction Kit (PE0230, Sigma-Aldrich, MO, USA) from 4-week old rosette leaves. Total protein concentration in the extracts was measured by carrying out a modified Lowry Assay, and denaturing polyacrylamide gel electrophoresis was performed to determine the purity and quality of the extracted protein. For quantification, equal amounts of total protein from each sample were loaded onto an automated capillary-based size western blotting system (ProteinSimple Wes System, San Jose CA, USA). Rubisco activase in each protein sample was detected using antibodies raised against rubisco activase (rabbit, AS10700; Agrisera, Sweden). Data analysis was carried out using Compass Software (ProteinSimple, San Jose CA).

### Measurement of 2-Phosphoglycolate

Eight-week-old plants were treated with 125 or 1000 µmol m^−2^ s^−1^ light for six hours. After treatment, leaf tissue was harvested and immediately frozen in liquid nitrogen before being stored at −80 °C. Samples were finely ground using a chilled mortar and pestle. Metabolites were extracted with 3:71:26 ethanol/formic acid/water (v/v). After centrifuging the supernatant was freeze dried, re-suspended in acetonitrile, and filtered through Mini-Uniprep Syringless Filter Devices, 0.2 µm (GE Healthcare).

LC/MS-MS was carried out on an Aquity UPLC Performance LC (Waters) and Quattro Premier XE (Micromass) in electrospray negative ion mode. The column used was an Acquity UPLC BEH Amide 1.7 µm, 2.1 x 100 mm (Waters). Mass Lynx (v. 4.1, Waters) was used for data acquisition and analysis. Acquity Binary Solvent Manager (v. 1.40.1248) and Acquity Sample Manager (v. 1.40.2532) was used as the binary solvent manager. Capillary voltage was 2.75 kV, source temperature was 120 °C and desolvation temperature was 350°C. Desolvation gas flow was set to 800 L/hr and collision gas flow was 0.15 mL/min. Selected reaction monitoring was used to quantify levels of 2-PG. The parameters for monitoring 2-PG were trace of 153 > 97 (m/z), cone voltage of 19 V, and collision energy of 17 V. Parameters for the internal standard (^13^C_6_ fructose 1,6-bisphosphate, Omicron) were trace of 345 > 97 (m/z), cone voltage of 16 V, and collision energy of 25 V. A multi-step gradient was used with 10 mM ammonium acetate in water (A) and acetonitrile (B). The flow rate was set to 0.3 mL min-1 throughout the entire run. 0-1.0 min, 5% A; 1.0-1.5 min, 5-100% A; 1.5-3.0 min, 100% A; 3.0-3.5 min, 100-5% A; 3.5-4.0 min, 5% A.

### Measurement of GAP, DHAP and TPI

About 500 mg fr wt of leaves were immediately freeze-clamped using aluminum blocks cooled on dry ice to stop metabolism fast enough to measure triose phosphates. Samples were weighed then fully pulverized using a Retsch Mill M300 (Retsch, http://www.retsch.com/). Cold 3.5% perchloric acid (2 µL per mg tissue), was added and tubes were placed on ice for 5 min incubation. Extracts were centrifuged at maximum speed at 4°C for 10 min. Approx. 500 µL of supernatant was recovered. Neutralizing buffer (2M KOH, 150 mM Hepes and 10 mM KCl), in the ratio of 0.25µL per µL of recovered supernatant, was added to the supernatant to bring the pH to ∼7. pH sticks were used to check the pH; the volume of neutralizing buffer was adjusted accordingly, and the volume of neutralizing buffer was recorded. Samples were frozen and thawed to precipitate salts, and centrifuged at maximum speed for 2 min. The supernatant was pipetted off for immediate GAP and DHAP assays, or frozen at −80 °C for future assays.

The amount of GAP and DHAP was measured using a dual wavelength filter photometer (Sigma ZFP22). Supernatant (50 µL) was added to 800 µL reaction buffer (100 mM Hepes buffer pH 7.6, 1 mM DTT, 1 mM KH_2_AsO_4_, 50 mM NAD and 50 mM ADP) in a cuvette, which was inserted into the cuvette holder in the spectrophotometer. Glyceraldehyde 3-phosphate dehydrogenase (5 U) (Sigma G-5537) was added to the cuvette and immediately mixed using a clean plastic stick. The difference of absorbance between the baseline and the maximum level was measured to estimate the level of GAP. Next, 5 U of triose phosphate isomerase (Sigma T-6285) was added to convert DHAP to GAP. We used A_334_ – A_405_ and an extinction coefficient of 6190 M^−1^ cm^−1^.

Triose phosphate isomerase activity was measured by linking the production of DHAP from GAP to oxidation of NADH to NAD through glycerophosphate dehydrogenase as described in Anderson (1971). TPI from crude protein extracts of Arabidopsis leaves harvested from four-week-old rosettes was assayed in the presence of varying concentrations of GAP (0, 0.2, 0.4, 0.7, 2 mM) and 2-PG (0, 10, 20, 40, 80 µM). The same dual wavelength filter photometer mentioned above was used to record the rate of change in absorbance at 334 - 405 nm.

### Quantitative RT-PCR

Arabidopsis total RNA was isolated using a Plant RNeasy kit according to the manufacturer’s instructions (Qiagen, https://www.qiagen.com), and treated with DNase I. 500 ng of each RNA sample was used for cDNA synthesis with random primers and iScript^TM^ cDNA Synthesis Kit (Bio-rad, http://www.bio-rad.com/). Quantitative real-time PCR was performed using a 7500 Fast Real-Time PCR System with Fast SYBR Green Master Mix (Applied Biosystems, http://www.appliedbiosystems.com). Gene expression was normalized to actin. Expression was determined in triplicate biological measurements.

### Statistics and Box Plots

Differences between WT and treatments were tested by one way ANOVA followed by Tukey’s test in Microcal Origin 8.0. Three levels of significance were test and indicated by +, *α* = 0.1; *, *α* = 0.05; **, *α* = 0.01. Box plots are presented with the box encompassing the middle two quartiles, the mean shown as an open square inside the box, the median as a line inside the box, and the whiskers showing the standard deviation of the data.

## Acknowledgements

We wish to thank Jim Klug and Cody Keilen (Growth Chamber Facility) of Michigan State University for their assistance. ABRC for seeds and Andreas Weber for *plgg1* seeds.

